# Does domestication trade-off stress tolerance for leaf growth? A search for evidence across eight Pooideae grass species

**DOI:** 10.1101/2024.11.30.626197

**Authors:** Jie Yun, Chenyang Yuan, Katherine Irelan, MJ Kabongo, Eldar Urkumbayev, David L. Des Marais

## Abstract

Plant domestication is thought to create trade-offs between high yield and stress tolerance, raising concerns about yield stability in future climates. Previous studies have found limited direct evidence for such trade-offs, often focusing on weakened defenses associated with higher growth rates. However, trade-offs can also occur when traits (such as yield in agriculture) optimized for favorable conditions perform less efficiently in stressful conditions. Deciphering the mechanisms driving these trade-offs is crucial for maintaining yield in changing environments. We examine leaf growth, a key trait influencing carbon assimilation, in eight species of grasses. We use a machine learning pipeline to automatically extract cell dimensions and positions from leaf microscope images to study cell kinematics, finding that domesticated plants generally have longer leaves, larger division zones and higher cell production rates. We found no clear evidence of trade-off between domestication and drought response in final leaf length. However, a trade-off is observed in development as wild species exhibited a smaller decrease in elongation zone size under drought than their domesticated counterparts. These nuanced trade-offs associated with domestication highlight the importance of examining physiological traits and mechanisms in greater detail, possibly informing breeding strategies to enhance crop resilience in the face of climate change.

**Highlight:** This study uses a high throughput pipeline to characterize leaf elongation responding to drought stress across eight species including barley, wheat, oat and wild relatives.

## Introduction

Plant domestication is associated with phenotypic change in many traits, including seed size, plant size, biomass ratio of harvestable tissues, and the synchronization of tillering and maturity (Meyer and Purugganan, 2013; Milla, 2023; Milla et al., 2024; Preece et al., 2017). One interpretation of these observations is that artificial selection for domestication changes the allocation of limiting resources, rather than directly targeting the relative growth rates of specific tissues (Evans and Evans, 1996; Simpson et al., 2017). Conceptualizing domestication as altered resource allocation suggests that there is an inevitable trade-off between plant life history components. This is because traits not experiencing direct selection, such as defense or stress response, may be negatively affected due to the relatively fewer resources they receive (Koziol et al., 2012; Mayrose et al., 2011). Indeed, current evidence supports the hypothesis that selection for larger seeds during domestication led to cascading effects such as larger plant size and faster growth rate needed to support high yield in agronomic settings (Milla and Matesanz, 2017; Preece et al., 2017).

The possible trade-off between high yield and reduced stress tolerance raises concerns about yield stability in future climates. Previous studies have sought to identify these trade-offs but have found limited direct evidence, often focusing on testing for weakened defense against pathogens or herbivores associated with higher plant growth rates (Simpson et al., 2017; Turcotte et al., 2014). Whether the paucity of unambiguous trade-offs between life history components in domesticated plants reflects a true absence of such trade-offs or merely an absence of evidence is unclear.

Under natural, resource-replete settings, there is evidence that traits associated with high resource acquisition will be favored by selection (Milla, 2023; Milla et al., 2015, 2014). These include more seminal roots (Golan et al., 2018), larger plant size and leaf area index (Milla et al., 2014; Milla and Matesanz, 2017), and higher specific leaf area (SLA) and maximum capacity of carboxylation (Gómez-Fernández et al., 2022). Trade-offs can occur when a trait selected under favorable environmental conditions performs less well in stressful conditions, as is often the case in agriculture, where yield is selected under optimal conditions (Agrawal et al., 2010; Agrawal, 2020). Leaf growth is a coordinated process comprising cell division and cell elongation (Fiorani et al., 2000). The leaves of domesticated grain species typically have larger area and relatively less dry biomass per area (i.e. have higher specific leaf area, SLA) as compared to their undomesticated relatives (Gómez-Fernández et al., 2022; Milla and Matesanz, 2017).

Because leaf anatomy, and by extension, patterns of leaf development, is a strong determinant of growth rate (Poorter, 1989; White et al., 2016), leaf growth parameters may have been an indirect target of artificial selection for high harvest index during domestication. However, the effects of domestication on mechanisms of leaf growth are not well understood. In this study, we focus on domestication effects on leaf elongation, adapting a kinematic approach to study the division and elongation dynamics of individual cells (Erickson and Silk, 1980). Drought stress is known to typically reduce leaf elongation rates and durations, decrease cell production, and limit cell expansion, although these responses can vary significantly between species (Lu and Neumann, 1998; Schuppler et al., 1998; Tardieu et al., 2000; Verelst et al., 2013). Despite this variability, no systematic and consistent analysis has been conducted to explore these effects comprehensively across multiple species. Here, we model leaf growth under both well-watered and soil drying conditions to assess what trade-offs, if any, may be present in domesticated grain species. To accomplish these aims, we developed a machine learning-based pipeline capable of automatically extracting cell dimensions and positions from leaf microscope images. This new tool offers a powerful approach to measure cellular dynamics in a range of plant species.

## Methods and Materials

### Plant growth and drought treatment

Focal species studied are listed in Table 1. Both domesticated wheat varieties (*Triticum turgidum* L. var. Durum ‘Langdon’ and ‘Svevo’, *Triticum aestivum* ‘Chinese Spring’) and wild emmer wheat (*Triticum turgidum* L. subsp. *dicoccoides* ‘Zavitan’ and ‘Vavilovii’) were subjected to a pre-germination treatment at 32°C dry conditions for 7 days, followed by 4°C dry conditions for 2 days. Domesticated oat (*Avena sativa* cv. ‘Sang’), wild oat (*Avena insularis* ‘BYU 207’), domesticated barley (*Hordeum vulgare* ‘Morex’), wild barley (*Hordeum vulgare* L. subsp. *spontaneum*), and *Brachypodium distachyon* (Bd21) seeds were placed at 4°C in water for 7 days to synchronize germination. After cold treatment, all seeds were soaked in Gibberellic Acid for 1 day to promote uniform germination. Seeds were then planted in Kord square pots (the HC Companies, OH, USA) filled with 70g of Profile porous ceramic rooting media (Profile Products, Buffalo Grove, IL).

**Table 1.** Species used in this study.

| Common name | Scientific name | Accession identifier | Name symbol | Domestication status | ploidy |
| --- | --- | --- | --- | --- | --- |
| wheat (wild emmer) | *Triticum turgidum ssp. dicoccoides* | Zavitan | Wheat-Z-Wild | Wild | 4x |
| wheat (bread wheat) | *Triticum aestivum* | Chinese spring | Wheat-CS-Domesticated | Domesticated | 6x |
| wheat (durum wheat) | *Triticum turgidum L. var. durum* | Langdon | Wheat-L-Domesticated | Domesticated | 4x |
| wheat (durum wheat) | *Triticum turgidum* L.<br>*var. durum* | Svevo* | Wheat-S-Domesticated | Domesticated | 4x |
| wheat (wild emmer) | *Triticum turgidum* L.<br>*subsp. dicoccoides* | Vavilovii*<br>(GRIN:PI 352322) | Wheat-V-Wild | Wild | 4x |
| oat | *Avena sativa* L. | Sang<br>(NPGS:PI386171) | Oat- Domesticated | Domesticated | 6x |
| oat | *Avena insularis* | BYU 207 | Oat-Wild | Wild | 4x |
| barley | *Hordeum vulgare* L.<br>*subsp. spontaneus* | (GRIN:PI 662181) | Barley-Wild | Wild | 2x |
| barley | *Hordeum vulgare* | Morex | Barley-Domesticated | Domesticated | 2x |
| purple false brome | *Brachypodium distachyon* | Bd21 | Bd21 | Wild | 2x |

\*The two species are only used for leaf elongation in this study, while the rest are used for both leaf elongation and cell elongation.

Before planting, the dry weight (DW) of each pot and its field capacity (FC) for water were recorded. FC was determined by saturating the pot with water and allowing it to drain overnight under gravity, with the weight difference between the saturated and dry pot used as the basis for subsequent soil dry-down control. The plants were grown in growth chambers (Biochambers FXC-9) with controlled conditions: 60% humidity, 500 µmol photons m^-2^ s^-1^ light intensity, and a temperature regime of 25°C during the day and 20°C at night (16h/8h day/night cycle). Initially, the plants were bottom watered every other day with tap water (pH 5-6), supplemented with DYNA-GRO GROW Liquid Plant Food 7-9-5 (Dyna-Gro, Richmond, CA, USA) until the appearance of the second leaf. From the appearance of the second leaf stage, control plants continued to be watered nightly without fertilizer, while drought treatment plants were subjected to controlled soil dry-down. Soil water content in the drought treatment group was reduced to 45% FC, and then water content maintained between 45-55% by watering at 10 a.m., 4 p.m., and 10 p.m. daily. These water levels were determined based on a pilot experiment showing that leaf hydraulic potential significantly decreases at this threshold (see Figure S1).

The third leaf was regularly checked at each watering. For both control and drought-treated groups, five replicates were used to measure the entire process of the third leaf’s elongation, and an additional five replicates were harvested on the fifth day after leaf appearance (approximately at the steady state of elongation) for microscopy.

### Leaf elongation measurements and model fitting for whole leaves

The third leaf of five plants were photographed every four hours using a Raspberry Pi camera. Leaf length was measured from the leaf base to tip using ImageJ, with a length scale correction applied based on a predefined standard to adjust for the perspective of each image. Elongation data for each sample was then used to fit a Beta-sigmoid model previously demonstrated to provide accurate fitting of grass leaf growth (Voorend et al., 2014). The model achieved an R² value greater than 0.95, indicating excellent fit (Figure S2). The model fitting yields parameters including final leaf length, the duration of leaf elongation (defined as the time between 10% and 90% of total leaf length), and the maximum elongation rate (representing steady-state elongation). These parameters enable statistical comparisons across species and treatments, with five replicates per species per treatment. For visualization, the fitted models are synchronized using cross-correlation, and the mean and 95% confidence intervals for the model fits of replicates are plotted over time for each group. The relationship between leaf length and time is modeled by the following equation:

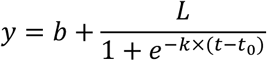

Where *t* is the thermal time of measurement, and *y* is leaf length. *b*, *L*, *k*, *t*_0_ are parameters from the model fitting.

Key parameters are

1. The **mature leaf length** is the maximum value of the function:

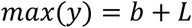
2. The derivative of the function represents the leaf elongation rate. The **maximum elongation rate occurs** (or steady state elongation rate) when *t* = *t*_0_

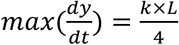
3. The **duration of elongation** is defined as the time between reaching 10% and 90% of the maximum leaf length. This duration is calculated as *l*_90_-*l*_10_ where

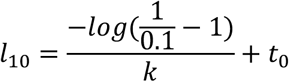

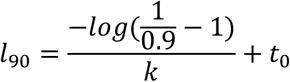

### Leaf cell length measurement

We used a second set of plants for leaf cell measurements. We carefully unwrapped an emergent third leaf from older leaves, and dissected as close as possible to its connection to the seed. This is done on the fifth day after the appearance of the third leaf, which corresponds to a steady state of leaf elongation as observed in the whole-leaf measurements, above. At this stage, the ligule is within 1mm of the leaf base. Dental impression material (Defend Impression Material, IL, USA) was applied to the base region (for leaves > 50mm) or whole leaf (<50mm) of the abaxial surface to create an imprint of the leaf. Thereafter, a thin layer of nail polish was applied to the imprint, then dried, and placed onto a microscope slide.

Microscopy images of imprints are taken with Zeiss Axiolab 5 with Axiocam 208 CCD camera (Carl Zeiss AG, Germany) with 10x magnification. A sequence of images were taken from the base of the leaf towards the tip with >20% overlap between images. To be consistent across different species, the distal border of the leaf division zone is diagnosed as the distal extent of symmetric division of stomata (McKown and Bergmann, 2020). (Note that this border is the same for trichomes, as the end of symmetric division of trichomes and stomata always co-occur based on our observations.) In this study, we focus on characterizing undifferentiated cells and sister cell elongation (after differentiation) along the leaf axis.

We developed a customized pipeline based on Cellpose (Pachitariu and Stringer, 2022) to perform automatic cell segmentation using machine learning. The pipeline is briefly described below; more details can be found in the supplementary information. We first fine-tuned a pre-trained Cellpose model on a small number of manually-segmented examples, iteratively correcting cell segmentations on test images to improve accuracy. This model was then used to identify cells in the remaining images, outputting a binary mask for each cell. The positions and dimensions of the masks are extracted from each image to represent the identified cells. The relative location of each image along the leaf axis is reconstructed from the displacement of an image in relation to its two neighbors. We measure the average directionality of the cells in each image to correct for the curvature of the leaf (especially in meristem regions), approximating the leaf as a piecewise linear curve.

Given the large diversity of species and developmental stage of cells measured in this project, we divided the images into ten different groups and trained a different model for each group to increase accuracy. These groups are necessary because of cell morphology differences (i.e., stomata development or cell differentiation), large cell-size variation (we observed > five times difference in length) and the need of using surrogates to identify cells occluded by dense trichomes. Of all the species considered, Oat proved particularly challenging, as we need to use the location of stomata cells to identify sister cells. These groups have overlaps to allow gradual changes, and the same sets of models are used in all the samples in each crop type to allow for comparison.

### Leaf cell length model fitting

Similar to leaf elongation, cell length along the leaf axis, beyond the division zone, can be modeled using a Beta-sigmoid function (Voorend et al., 2014). A custom Python script was used to fit this model, where the distance from the base of the leaf serves as the independent variable (*x*) and cell length as the dependent variable (*y*). The model fitting achieved an R^2^ value higher than 0.75 in all samples, with an average R^2^ of 0.92 (Figure S3). The model fitting allows us to estimate key parameters such as mature cell length, the number of cells in a mature leaf, cell flux, and metrics related to both cell division and elongation. These include division zone size, cell number in the division zone, average cell size in the division zone, average division rate, elongation zone size, elongation zone cell number, elongation duration, and the average relative elongation rate.

Specifically, the model is defined as:

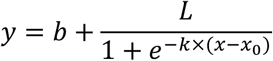

Where *x* is the location with reference to end of division zone, and *y* is cell length. *b*, *L*, *k*, *x*_0_ are parameters from the model fitting. Parameters are modified from (Kavanová et al. 2006).

Key parameters are:

1. The **mature cell length** (*L_f_*) is given by:

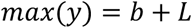
2. **Number of cells in a mature leaf** is calculated as the ratio of mature leaf length and mature cell size. Mature leaf length is determined from the leaf elongation measurements.
3. **Cell Flux** (*F*, cells °Cd^-1^) or cell production rate, the rate that cells are displaced at a particular position, is estimated from the leaf elongation rate (*LER*) from the leaf elongation measurements and mature cell length (*L_f_*)

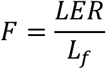
4. **Elongation zone size** (*Lge*) represents where the cell achieves 95% of their mature cell size.

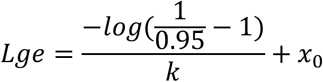
5. Elongation zone cell number **(*N_e_*)**

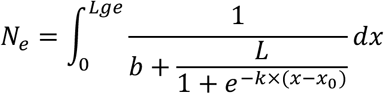
6. Cell displacement velocity at a certain location in the elongation zone (*v*(*x*)) is composed of the elongation of all the cells located between the cell and the base. It can thus be approximated as proportional to the local cell length. The velocity of the cell at the end of the elongation zone equals the leaf elongation rate (*LER*).

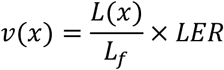 **Average relative elongation rate in elongation zone** 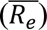 is the calculated as the average of displacement velocity at beginning of the elongation zone (end of division zone) (*v_d_*) and the end of the elongation ( *v_e_*), normalized by elongation zone length

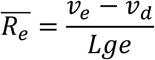
7. To convert the location to the time needed from the beginning of the elongation zone, number of cells between location *x* and the beginning of the elongation zone (*N_e_*(*x*)) is connected to time through flux rate

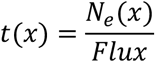 **The average elongation duration** (*T_e_*) is then calculated as

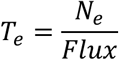
8. **Number of cells in division zone** (*N_div_*) is directly calculated based on the length of the division zone and the average of division zone cell size.
9. **Average division rate** (*D*, cell cell^-1^ h^-1^) is derived from cell production rate to number of cells in division zone:

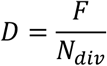

This model allows us to precisely quantify leaf growth dynamics and cell division and elongation processes, which are key to understanding how different conditions, such as drought or domestication, affect plant development.

### Statistical analysis

All statistical analyses were performed in R (R Core Team, 2022). The parameters calculated above were fitted into linear models (*lm*) or linear mixed-effects models (*lmer*) (Bates et al., 2015). Akaike Information Criterion (AIC) was used to select the best models, with crop, treatment, and domestication as factors and accessions (in wheat) and batch included as random factors, when appropriate. For parameters derived from two datasets (e.g., final cell number, the ratio of final leaf size to mature cell size), estimates were obtained by bootstrapping each parameter 2000 times within groups and matching them randomly to study treatment or domestication effect (results shown in the heatmaps). For models that included interaction terms (Table 2), the *anova* function from the *car* package (Fox et al., 2023) was used to compute p-values. The *emmeans* package (Lenth et al., 2024) was employed to calculate the marginal effects of main factors. *Brachypodium*, lacking a domesticated counterpart, was excluded from analyses for effect size involving balanced models. However, its effects were analyzed separately, with all other parameters treated as random factors.

**Table 2.** ANOVA of domestication, treatment and crop effect p-value.

|  | leaf<br>length | leaf<br>elongation<br>duration | leaf<br>elongation<br>rate | mature<br>cell size | division<br>zone<br>size | division<br>zone cell<br>size | division<br>zone<br>cell<br>number | elongatio<br>n zone<br>size | elongation<br>zone cell<br>number |
| --- | --- | --- | --- | --- | --- | --- | --- | --- | --- |
| crop | 2.7E-27 | 6.5E-13 | 1.6E-05 | 0.0E+00 | 9.7E-15 | 1.3E-63 | 4.0E-15 | 2.0E-05 | 1.9E-29 |
| Treatment | 6.4E-45 | 9.7E-21 | 1.7E-46 | 2.8E-17 | 3.7E-02 | 7.9E-01 | 6.9E-02 | 2.2E-33 | 7.7E-32 |
| domestication | 7.8E-09 | 3.1E-06 | 6.4E-23 | 2.9E-07 | 5.9E-07 | 1.6E-01 | 5.4E-06 | 2.9E-08 | 2.5E-04 |
| crop:<br>Treatment | 6.4E-02 | 2.1E-04 | 2.6E-01 | 2.1E-02 | 4.7E-02 | 9.6E-02 | 4.8E-02 | 1.0E-04 | 9.6E-08 |
| crop:<br>domestication | 3.0E-01 | 1.9E-01 | 1.6E-04 | 5.3E-11 | 3.7E-01 | 6.6E-02 | 4.3E-01 | 1.4E-01 | 2.3E-01 |
| Treatment:<br>domestication | 2.4E-01 | 3.7E-01 | 5.5E-03 | 5.2E-02 | 9.1E-01 | 1.8E-02 | 7.8E-01 | 8.4E-06 | 1.6E-02 |
| Crop:<br>Treatment:<br>domestication | 1.7E-01 | 1.2E-02 | 8.6E-02 | 2.4E-02 | 4.2E-01 | 4.9E-02 | 9.0E-01 | 1.2E-01 | 6.1E-02 |

For pairwise contrasts, p-values and response proportions were extracted from regression summaries of one main factor. These results were then visualized as heatmaps in *ggplot* (Kassambara, 2023; Wickham, 2016), with p-values indicated by asterisks and response proportions represented on a color scale. To visualize trends in leaf length over time, and cell length along both time and distance from the leaf base, fitted models for each sample were plotted using *ggplot*, with the mean and 95% confidence intervals aggregated within each group.

## Results

### Domestication increases mature leaf length and the rate of leaf elongation

Mature leaf length varied from 38mm to 232mm among species and treatments in our study, with the smallest length observed in *Brachypodium* under drought stress and largest length observed in domesticated wheat (Chinese spring) under the control condition. Generally, different crops have very different leaf lengths (p-value = 2.7E-27, Table 2) with *Brachypodium* having the smallest leaves (Table S1).

Both final leaf length (p-value=7.8E-09, Table 2 and Figure 1A) and the rate of leaf elongation (p-value=6.4E-23, Table 2 and Figure 1A) are greater in domesticated plants as compared to their wild relatives, with different magnitudes across crops (interaction term p-value=1.6E-04, Table 2). Elongation rate is the combined effects of cell production rate and final cell size. In our experiments domesticated plants show a universal increase of cell production rate, defined as cell flux, calculated as the ratio of leaf steady state elongation rate and mature cell length (Figure 1D). To investigate the possible mechanism of this domestication effect, we observe that the division zone size always increases in a domesticated species as compared to its wild relative (p-value=5.9E-07, Table 2 and Figure 1C, D), as does the number of cells in the leaf division zone (p-value=5.40E-06, Table 2 and Figure 1D). Domestication effects on final cell size varied among crops (interaction term p-value=5.30E-11, Table 2 and Figure 1D). However, domestication in general is found to increase both elongation zone size (p-value=2.90E-08, Figure 1B,D) and elongation zone cell number (p-value=2.50E-04, Figure 1D). Note, however, that the comparison between wheat (Chinese spring) and wild wheat shows an opposite trend. These results are consistent with the hypothesis that domestication leads to higher elongation rate and, ultimately, leaf length.

**Figure 1.**
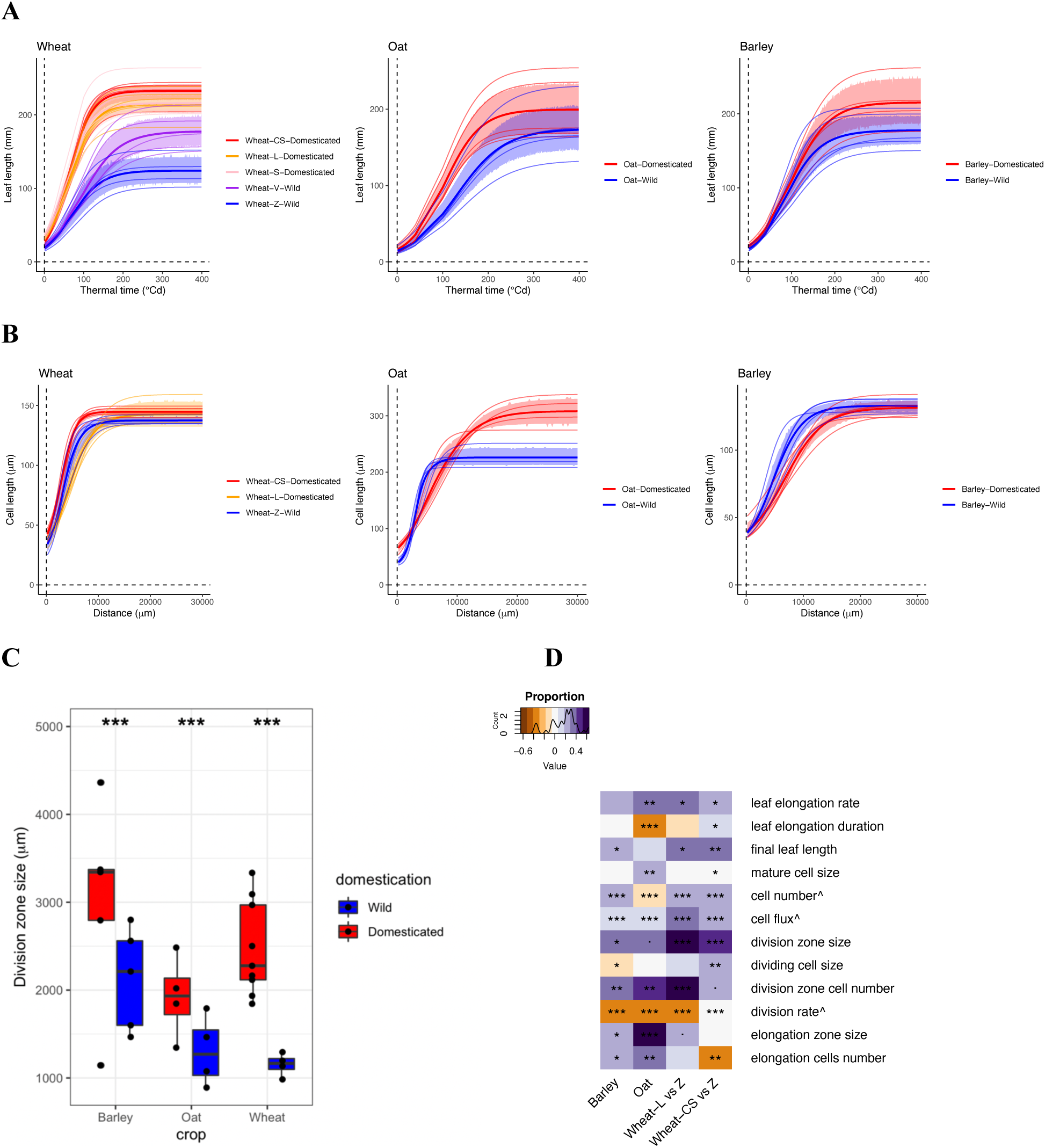
Leaf growth comparison between domesticated and wild species in barley, oat and wheat when grown under well-watered condition. **A)** Leaf elongation as a function of degree days. **B)** Lengths of individual epidermal cells along the leaf axis. **C)** Size of leaf division zone. **D)** Heatmap summarizing the statistical analysis of domestication effects on leaf growth parameters derived from **A, B** and **C**. Colors represent the direction and magnitude of changes (+ domestication positively affects the value, - domestication negatively affects the value), with asterisks denoting the significance levels of these changes (. p < 0.1, * p < 0.05, ** p < 0.01, *** p < 0.001). ^ parameters are calculated as ratio based on permutations of two datasets

Leaf elongation duration can compensate for changes in leaf elongation rate to some degree. Leaf elongation – measured in thermal time units – varied, with the lowest values observed in Bd21 under control conditions (114.8 °Cd). Crop species have different elongation durations (p-value= 6.5E-13, Table 2). Generally, *Brachypodium* has the shortest leaf elongation duration, whereas wheat, oat and barley have similarly long leaf elongation duration. Domestication decreases elongation duration by 34 °Cd (p-value =3.1E-06, Table 2 and Figure 1A). Note, however, this trend is driven by the contrast in oat, and the comparison between wheat (Chinese spring) and wild wheat shows an opposite trend.

### Drought decreases mature leaf length and changes the rate and duration of elongation

Final leaf length decreases in all sampled species under drought conditions (p-value= 6.4E-45, Table 2, Figure 2A, D). This is consistent with decreasing elongation rate under drought (p-value=1.7E-46, Table 2, Figure 2A, D), although increased elongation duration compensates for some of the possible reduction of mature length caused by soil drying (p-value=9.7E-21, Table 2, Figure 2A, D). The decreased elongation rate is a result of both the decreased cell flux (Figure 2D) and final cell size under soil drying (p-value=2.80E-17, Table 2, Figure 2B, D). Wild oat is an exception to these broad trends, as soil drying does not decrease its cell size (Figure 2B, D). The decreased cell flux is related to decreased division zone size as well as cell number in oat and wheat (Figure 2C, D), thus the division rate, defined as the ratio of cell flux and division zone cell number, decreases except in wild wheat (Figure 2D).

**Figure 2.**
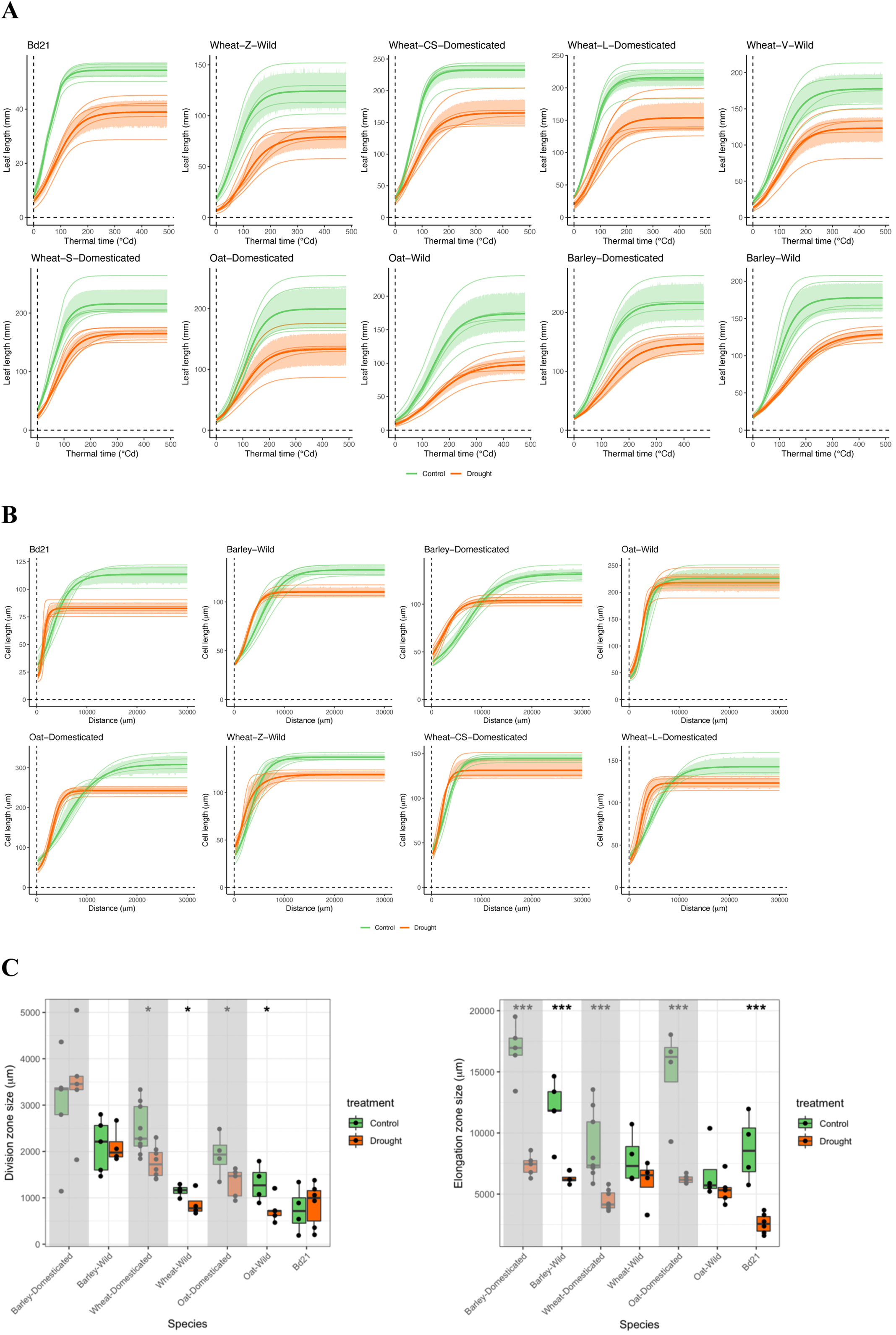
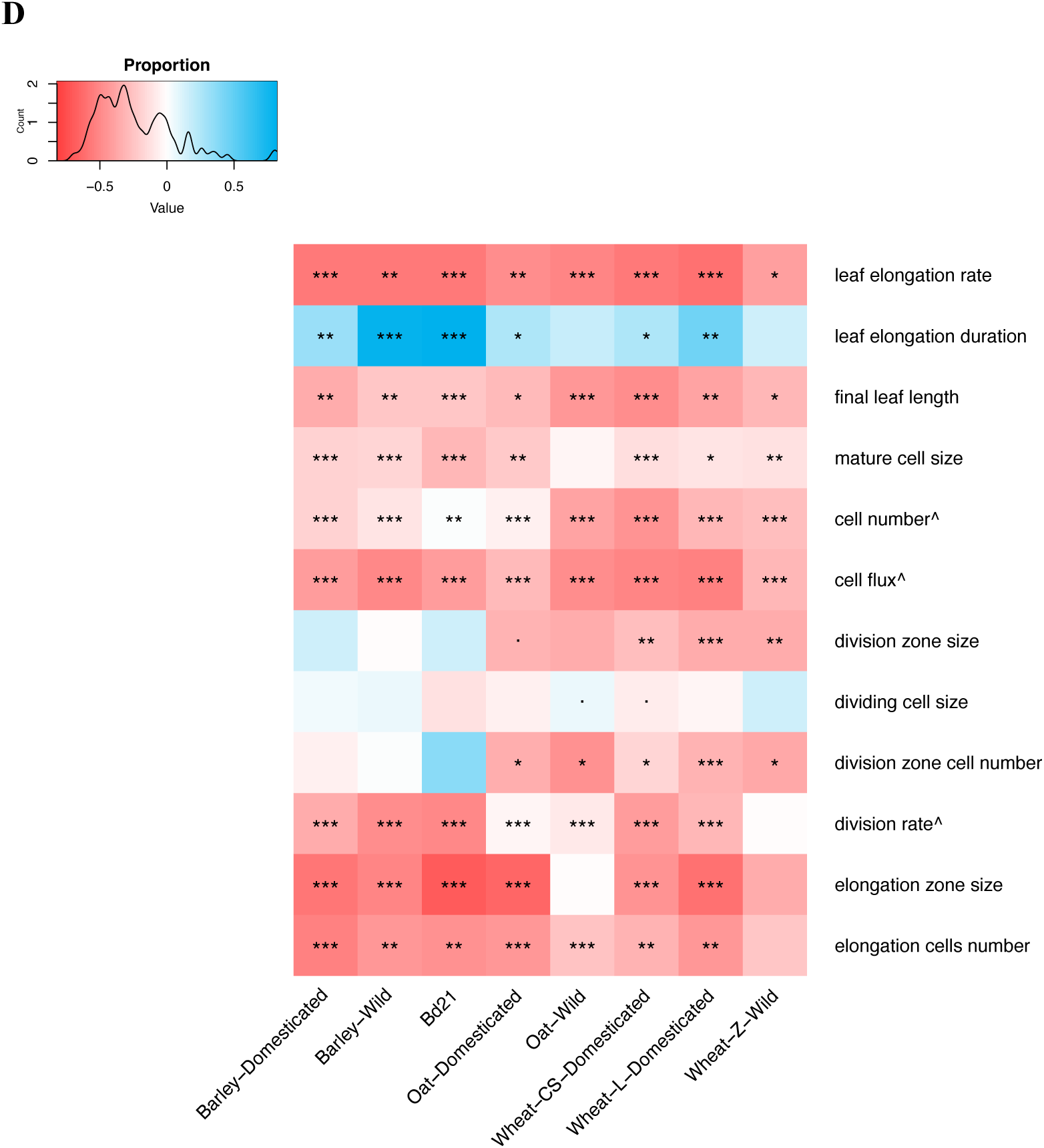
Leaf growth comparison between drought and well-watered plants across all the species. **A)** Comparison of leaf elongation along time between drought and well-watered plants **B)** Comparison of cell profile along leaf axis between drought and well-watered plants **C)** Comparison of leaf division zone size and elongation zone size under drought condition across species **D)** Heatmap summarizing the statistical analysis of drought effects on leaf growth parameters derived from **A, B** and **C**. Colors represent the direction and magnitude of changes (+ drought positively affects the value, - drought negatively affects the value), with asterisks denoting the significance levels of these changes (. p < 0.1, * p < 0.05, ** p < 0.01, *** p < 0.001). ^ parameters are calculated as ratio based on permutations of two datasets

Final cell size could be driven by the smaller elongation zone under drought (p-value=2.20E-33) which is observed in all species except wild oat (Figure 2C, D). This difference is also reflected in the fewer number of cells observed in the elongation zone under drought (p-value=7.70E-32; Figure 2D). Elongation zone and cell number show a significant interactive effect of treatment with species (p-value =1.00E-04 and 9.60E-08, respectively) suggesting considerable variation among species in their response; regardless, both traits are observed to decrease in most species under drought (Figure 2C, D). The elongation rate and duration generally show opposite effects of drought showing a compensation effect and diversity across species in the degree of compensation (Figure S4B, D). Thus, a consistently smaller elongation zone is observed in all plants under drought treatment and accordingly, smaller mature cells, again with the exception of wild oat. Interestingly, we observe that Bd21 has a constant cell number under drought stress (Figure 2D), consistent with prior results in this system (Verelst et al., 2013).

### Limited trade-off between domestication and growth responses to drought

Finally, we test the hypothesis that the positive effect of domestication on growth rate under control, well-watered, conditions results in reduced plasticity in growth rate of crop species under drought conditions. In domesticated plants, the reduction in final leaf length under drought conditions is similar to that seen in their wild relatives (Figure 2D), suggesting no trade-off exists between leaf growth in optimal and drought-stressed conditions (with interaction p-value= 2.4E-01, Table 2). However, contrasting patterns are observed in the cellular and developmental traits affecting final leaf length. Consistent with a larger division zone and greater number of dividing cells of domesticated plants as compared to their wild relatives under control conditions, domesticated plants maintained this trend under drought conditions (Figure 2C, D). This also suggests no interaction between domestication and drought effect on division zone size (p-value=9.1E-01, Table 2). We do observe a significant interaction between domestication and water treatment on elongation zone length (p-value=8.4E-06, Table 2), suggesting that wild and domesticated species respond to drought to differing degrees. Specifically, the elongation zone tends to be longer in domesticated plants under well-watered conditions but similar in size to wild species when exposed to soil drying (Figure 2C, D). This leads to the marginal trade-off effect in leaf elongation rate (p-value=5.5E-03, Table 2, Figure 2D) although to varying degrees among species. This result suggests that much of the gain in the growth capacity of domesticated plant leaves is realized under ideal conditions with no concomitant benefit – or penalty – under drought.

## Discussion

In this study, we systematically compared eight species, comprising four pairs of wild and domesticated species, to assess the effects of domestication on leaf growth under control and drought stress conditions. The model grass *Brachypodium distachyon* was also included, as prior work suggested that this wild species differs in its drought growth response as compared to domesticated grain crops (Verelst et al., 2013). We used a kinematic approach to investigate the roles of cell division and elongation using a high-throughput pipeline characterizing cell profiles at steady-state of elongation and leaf elongation dynamics.

Consistent with previous studies, we find that domestication generally results in longer leaves under well-watered conditions (Milla and Matesanz, 2017), supporting the hypothesis that domestication leads to more resource-acquisitive tissues and higher growth rate (Milla, 2023; Milla et al., 2014). Our study suggests these longer leaves can be explained by a higher leaf elongation rate, driven in part by increased cell flux. A key mechanism driving increased cell flux may be enlargement of the division zone and an increase in the number of cells within this zone. Previous research has hypothesized that, while overall growth rates increase during domestication, relative growth rates—when standardized to plant size—remain unchanged. Thus the increased absolute growth rates are possibly due to early seed vigor and cascading developmental effects (Milla and Matesanz, 2017; Preece et al., 2017). The increase in division zone size could be a contributing mechanism supporting this hypothesis.

Although there is no direct evidence connecting seedling vigor and division zone size, the division zone size is determined by hormone gradients: gibberellin levels spatially control cell division (Nelissen et al., 2018, 2012) and also affect seed germination (Vishal and Kumar, 2018). Many traits comprise the plant domestication syndrome (Alam and Purugganan, 2024); our results suggest that the size and activity of the leaf division zone may warrant further assessment in this context. This trait, which may have been unconsciously selected during domestication like many other traits (Purugganan, 2022), could provide insights into how domesticated plants optimize resource acquisition under favorable conditions.

Under drought conditions, leaf growth is inhibited either due to limited resources (e.g., sugar, turgor pressure) or via active regulatory signaling. While a prolonged elongation duration partly compensates for the decrease of leaf elongation rate, we found that drought typically leads to a shorter final leaf length, as found previously (Schuppler et al., 1998; Verelst et al., 2013). Mechanistically, cell size and cell number tend to decrease in most species in response to drought. This reduction in cell flux may stem from either a smaller division zone or from decrease in division rate. Previous studies have shown that meristem activity and cell cycle progression may be inhibited under drought stress (Schuppler et al., 1998; Skirycz and Inzé, 2010), directly affecting cell division. The reduction in cell size under drought can be attributed to a smaller elongation zone or to decrease in the number of elongation cells in this zone across species. Drought particularly impacts cell elongation, as cell expansion requires both cell wall loosening and turgor pressure, both of which might be compromised under water-limited conditions. Other factors, such as ABA signaling and sugar availability, likely also play a significant role in regulating cell elongation under drought stress (Skirycz and Inzé, 2010; Tardieu et al., 2010).

An intriguing exception to the general pattern of final cell number decrease under drought stress was *Brachypodium*, where the final cell number did not decrease under drought conditions, with the division zone size remaining unchanged. This finding aligns with previous studies (Verelst et al., 2013) and highlights *Brachypodium*’s potential ability to recover rapidly upon rewatering, perhaps as an adaptation to the rainfall regime of its native Mediterranean distribution.

The domestication trade-off hypothesis posits that artificial selection imposed under favorable conditions under stress resulted in reduced plasticity to environmental stress. However, our findings challenge this hypothesis: domesticated plants maintained their longer leaves under drought, consistent with previous studies investigating trade-offs between domestication and stress tolerance or defense response at higher-level physiological traits (Simpson et al., 2017; Turcotte et al., 2014). More broadly, the concept of a defense-growth rate trade-off has likewise been challenged (Kliebenstein, 2016). We further investigated potential trade-offs at the cellular level. Consistent with the higher-level findings, domesticated plants maintained their longer division zones and higher numbers of dividing cells under drought stress. However, the longer elongation zones observed in domesticated plants under control conditions were no longer significantly higher than wild species under drought, suggesting a potential trade-off in this aspect of leaf development.

We attribute the limited evidence supporting trade-offs to two potential factors. First, the absence of trade-offs at the physiological level may be specific to the traits measured in this study, particularly leaf elongation. Other aspects of leaf development, such as width, biomass, or chlorophyll content, may reveal different patterns. These traits, which were not the focus of our study, could play a role in carbon assimilation and overall plant fitness under stress. We observed a trade-off in elongation zone size which may warrant further investigation if it leads to increased fitness. Second, trade-offs may be more pronounced in simpler processes governed by fewer regulatory factors, whereas complex traits like overall leaf growth—which involve cell division, elongation, and growth duration, and may be controlled both by active regulatory processes as well as simple resource limitation—may dilute or compensate for trade-off effects. It is also possible that trade-offs are less common in domesticated plants than hypothesized. The long history and diversity of agricultural practices may have mitigated potential trade-offs by allowing for the selection of traits that maintain stable growth under both favorable and stressful conditions. For example, the enlargement of the division zone may represent a domestication trait that confers advantages regardless of environmental conditions. Moreover, the trade-off effects may be subtle, particularly when certain intrinsic genetic traits exert a strong influence, making them difficult to detect in the presence of large variances. However, if this is the case, the importance of identifying such trade-offs may be diminished, as these strong genetic influences could override potential trade-off impacts. Additionally, some patterns of trade-offs may be tied to specific agricultural practices or the evolutionary history of domestication. This is consistent with the finding that components underlying growth responses to domestication are found to be specific to crop types, phylogeny and climate (Gómez-Fernández et al., 2022).

Finally, we presented a high-throughput phenotyping pipeline to extract critical leaf elongation and cell division and expansion features from images and evaluate the significant effects of domestication and treatment effect. This approach substantially reduces the labor involved in manually measuring cell length using traditional methods. It is also less prone to systematic errors such as cell location shifts, which is mitigated through automated image stitching. Furthermore, our pipeline is versatile and can be readily applied to other fields, such as spatial single-cell omics analysis (Nobori et al., 2023), where cell outlines and locations are required. The high-throughput nature of our pipeline enables large-scale screening for phenotypes of interest, especially in the context of phylogenetic information, making it highly relevant in the era of abundant data and advanced genotyping technologies.

## Supplementary data

Supplementary Table 1. ANOVA of domestication, crop and treatment effect on growth parameters

Supplementary Figure 1. Hydraulic potentials change along with soil water content in different species

Supplementary Figure 2. Details of leaf elongation data and curve fitting

Supplementary Figure 3. Details of cell size data and curve fitting

Supplementary Figure 4. Cell size increase over time and two elongation parameters

Supplementary File 1. Details of high throughput pipeline for microscopy image analysis

Supplementary Data Set 1. Leaf elongation dataset

Supplementary Data Set 2. Leaf elongation model fitting

Supplementary Data Set 3. Cell size dataset

Supplementary Data Set 4. Cell size model fitting

## Author contribution

JY, DLD: conceptualization; JY, CY: methodology; JY, KI, EU, MK: data collection; JY: formal analysis; JY: writing - original draft; JY, CY, DLD: writing - review & editing; JY: visualization; DLD: supervision; DLD: funding acquisition

## Acknowledgement

We thank Mingcheng Luo, Steven Xu, Jeff Maughan, Gary Muehlbauer, Nils Stein and USDA GRIN for providing seeds of our focal species.

## Data availability

The leaf elongation package and tutorial is available at https://github.com/yuanchenyang/leaf_elongation

The code to regenerate all the results in this paper (including the parts using the above package) are available at https://github.com/jyjiey/leaf_elongation_comparison

The original microscopy images and leaf elongation data that support the findings of this study will be available at Dryad at Yun, Jie; Yuan, Chenyang; Irelan, Katherine et al. (Forthcoming 2024). Does domestication trade-off stress tolerance for leaf growth? A search for evidence across eight Pooideae grass species [Dataset]. Dryad. https://doi.org/10.5061/dryad.47d7wm3pp

## Abbreviations

CS: Chinese spring
V: Vavilovii
Z: Zavitan
S: Svevo
L: Langdon

## Supplementary information

**Supplementary Table 1.** ANOVA of domestication, treatment and crop marginal effect size.

|  | leaf<br>length | leaf<br>elongation<br>duration | leaf<br>elongation<br>rate | mature<br>cell size | division<br>zone<br>size | division<br>zone cell<br>size | division<br>zone<br>cell<br>number | elongatio<br>n zone<br>size | elongation<br>zone cell<br>number |
| --- | --- | --- | --- | --- | --- | --- | --- | --- | --- |
| ww | 200.29 | 178.47 | 1.29 | 184.86 | 2002.50 | 27.76 | 73.96 | 11208.27 | 130.33 |
| wd | 137.32 | 240.00 | 0.72 | 157.54 | 1712.41 | 28.30 | 62.46 | 5953.23 | 77.16 |
| W | 146.52 | 227.23 | 0.81 | 161.38 | 1386.52 | 27.88 | 51.88 | 7372.16 | 97.80 |
| D | 191.09 | 191.25 | 1.20 | 181.02 | 2328.39 | 28.18 | 84.54 | 9789.34 | 109.69 |
| barley | 194.94 | 217.73 | 1.16 | 119.01 | 2800.83 | 29.69 | 102.23 | 8592.12 | 126.80 |
| Bd21 | 90.10 | 120.74 | 0.79 | 106.02 | 1386.15 | 19.78 | 64.81 | 5118.19 | 92.52 |
| oat | 177.54 | 207.26 | 1.07 | 261.52 | 1428.33 | 35.50 | 44.38 | 6335.54 | 50.40 |
| wheat | 136.73 | 201.58 | 0.81 | 129.97 | 1622.26 | 24.41 | 67.63 | 6231.08 | 92.60 |

**Supplementary figure 1.**
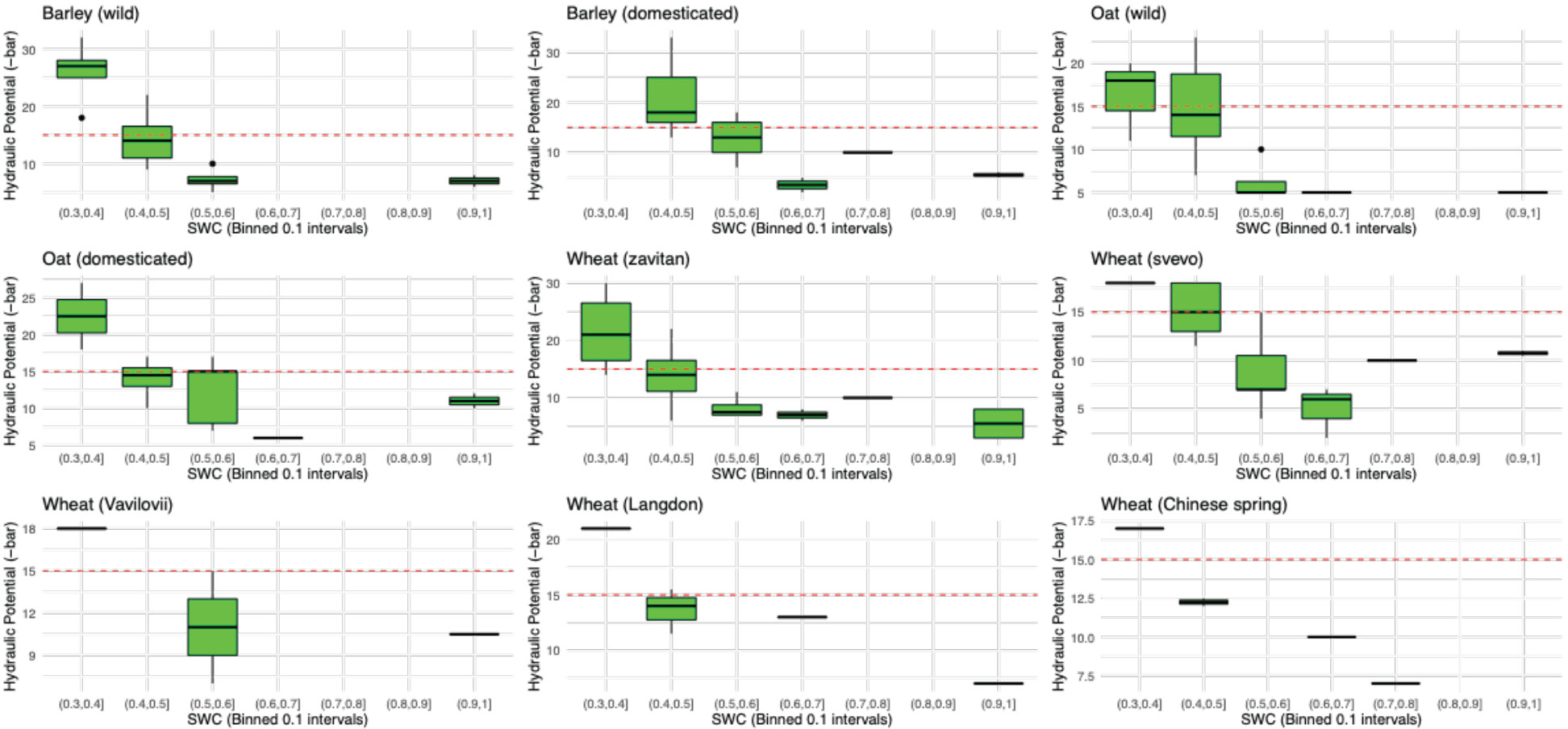
Hydraulic potential changes along soil water content (SWC) in each species. Red line shows where hydraulic potential reaches -15 bar. Leaf water potential was measured by the Scholander Pressure Chamber Model 670 (PMS Instrument Company, Albany, OR) on the youngest fully expanded leaf of sampled plants.

Supplementary figure 2 details of leaf elongation data and curve fitting (see supplementary files)

Supplementary figure 3 details of cell size data and curve fitting (see supplementary files)

**Supplementary Figure 4.**
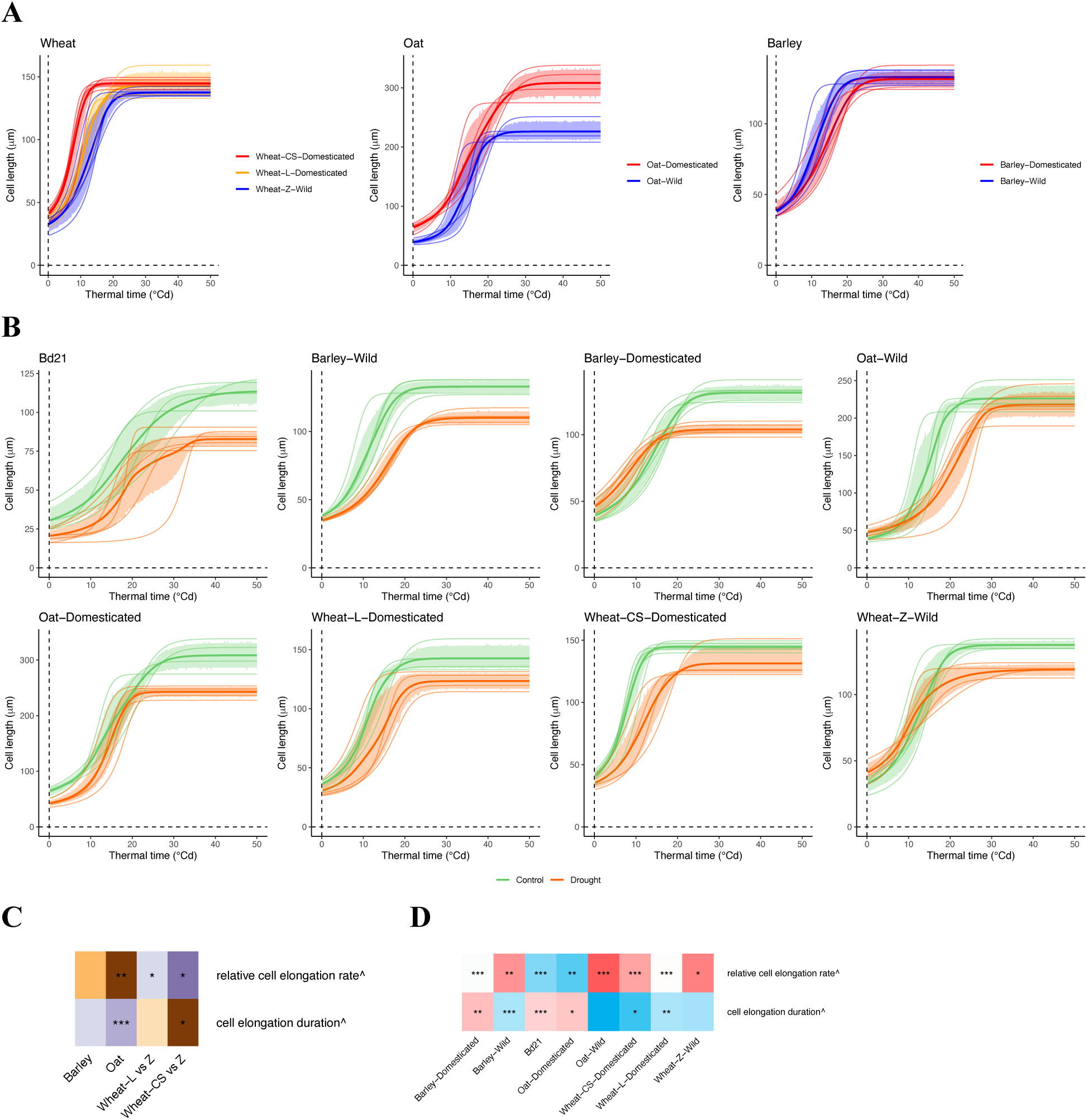
**A**) Comparison of cell profiles as a function of thermal time between wild and domesticated species in three crops grown under well-watered condition **B**) Comparison of cell profiles as a function of thermal time between control and drought in all species **C**) and **D**) Heatmap summarizing the statistical analysis of drought effects on leaf growth parameters derived from **A** and **B**. Colors represent the direction and magnitude of changes (Purple + domestication positively affects the value, Brown – domestication negatively affects the value in **C**; Skyblue + drought positively affects the value, Salmon – drought negatively affects the value in **D**), with asterisks denoting the significance levels of these changes (.<0.1, *<0.05, **<0.005, ***<0.0005). Both of the parameters are calculated as ratios from two datasets.

## Supplementary information File 1-- Microscopy image analysis

In this study, we developed a pipeline to process microscopy images for leaf cell length elongation measurements. This framework and its components were specifically designed for comparative analysis of different varieties or treatments; the parameters may be adjusted to segment anatomical structures to extract their dimensions and locations in an unbiased way. Our motivations were as follows:

1. Given a large number of samples collected (5666 images in total), it is labor intensive and also error prone to hand mark cells. We instead developed a machine-learning based cell segmentation pipeline using Cellpose (Pachitariu and Stringer, 2022). To do this, we fine-tuned a pretrained model on a few hand-labeled examples so the new model can be used on the remainder of images. In the end we identified 533977 regions of interest (ROI, including cells and other referred features, e.g., trichome) in total across all images.
2. This framework also increases flexibility of cell identification by straightforward adjustment or addition of functioning blocks. Given the large diversity of species in this project, we divided images based on different crops and stages of sister cells into 10 models where the same set of models are used by all the samples in each crop type to allow for comparison. Different models are chosen when there are differences in cell morphology (stomata development, or meristem cell vs sister cells) or cell size difference (>5 times) to increase accuracy, with overlaps between groups. In some situations, the direct measure of one type of cell is not possible. For example, *Brachypodium* and wheat have long trichomes, which cover most of the cells along the leaf (in *Brachypodium*) or elongation zone (in wheat). While most species have one cell file of stomata cells between two lines of sister cells, Oat has two cell files of stomata cells between two lines of sister cells. This generally misleads the identification of sister cells, which were the foci of the current study. When it is not possible to directly segment sister cells we identify other types of cells as references (i.e. using surrogates to identify cells occluded by dense trichomes or filtering sister cells by stomata cells in oat).
3. Accurate estimation of a cell’s location along the leaf axis is critical for calculating kinematics. Here we provide a framework to do so. Given a series of microscope images of a leaf, we identify the displacement of each image relative to its two neighbors to reconstruct the absolute distance of each image from the leaf base. We account for the curvature of the leaf when calculating the distance along the leaf axis by approximating a leaf as a piecewise linear curve.
4. Besides saving labor, another advantage of our method is to measure several files of cells to increase the accuracy. Given that cell lengths change gradually along the leaf axis, and each parallel cell file should change similarly along the leaf axis, the cells recognized spatially nearby can be used as replicates. This increases the accuracy of the trend-fitting along the leaf axis in the next step. Additionally, borrowing dimensional information from adjacent cell files also lowers the required image quality allowing for high throughput imaging collection.

The details of the pipeline follow. All components are in a package available at https://github.com/yuanchenyang/leaf_elongation

1. Images are taken on the abaxial surface along the leaf axis so that there is at least one image at any distance away from the base of the leaf in the beginning first 5cm or whole leaf if it is <5cm, which captures the division and elongation and part or all of the mature zone. Since the photos usually have more than one cell file of interest at the same distance to the base, they are considered replicates.
2. Given microscope images of a leaf, we need to identify the displacement of each image relative to its neighbors, to reconstruct the absolute distance of each image from the leaf base. One challenge is that consecutive images can be taken on different focal planes, so that their overlapping areas do not match exactly. We addressed this problem by developing a customized stitching algorithm. In brief, we first apply a filter that isolates high-, mid-, or low-frequency image features, then find the max 2D cross correlation across the filtered images using Fast Fourier Transforms (FFTs). The resulting algorithm (implemented in Python) achieves close to 100% accuracy, finding the correct displacements between two images even though they can have very different focus (see example in Fig).

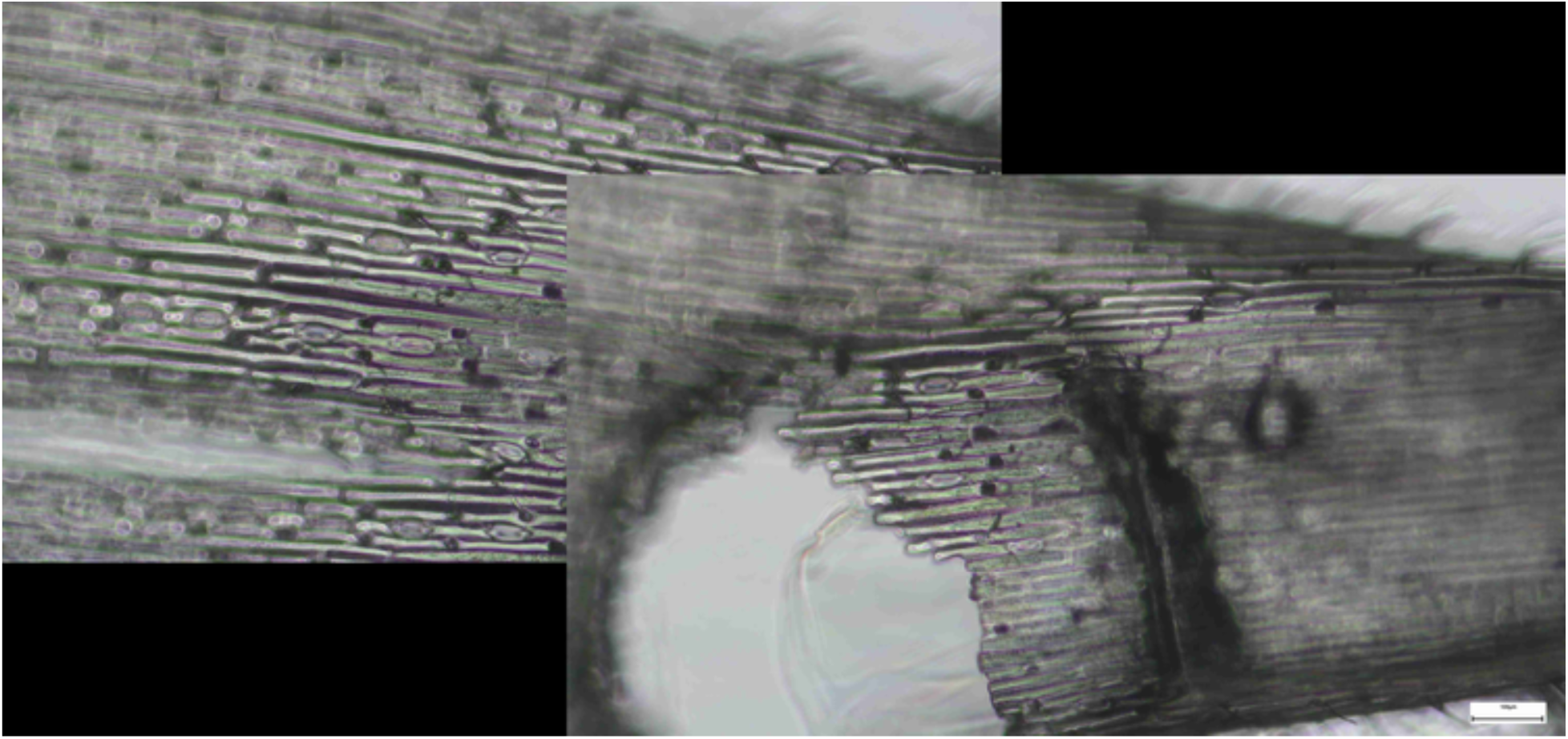
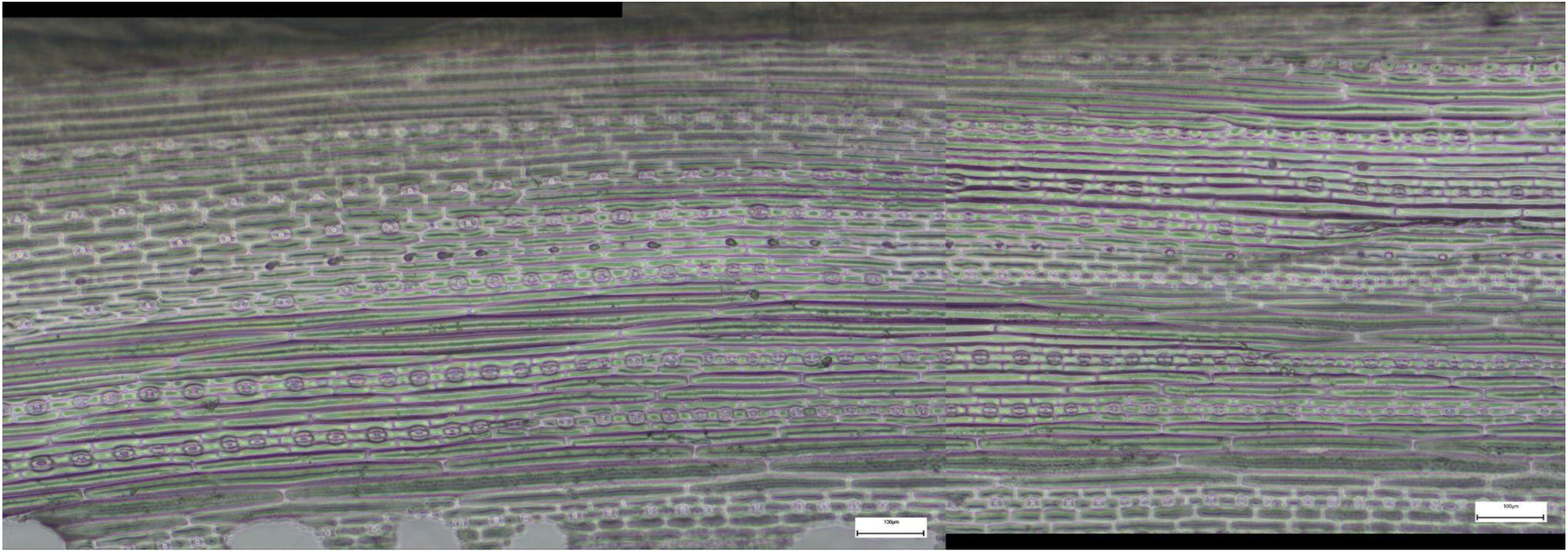
3. We also need to account for the curvature of the leaf when calculating the distance of each cell along the leaf axis. Hence we approximate a leaf as a piecewise linear curve, with each image having a constant direction, determined by the median orientation of all cells in that image. This is accomplished by a custom script that first computes the direction of edges with a Sobel filter, then fits a distribution to the directions found using Gaussian kernel density estimation, before returning the mode of this distribution.

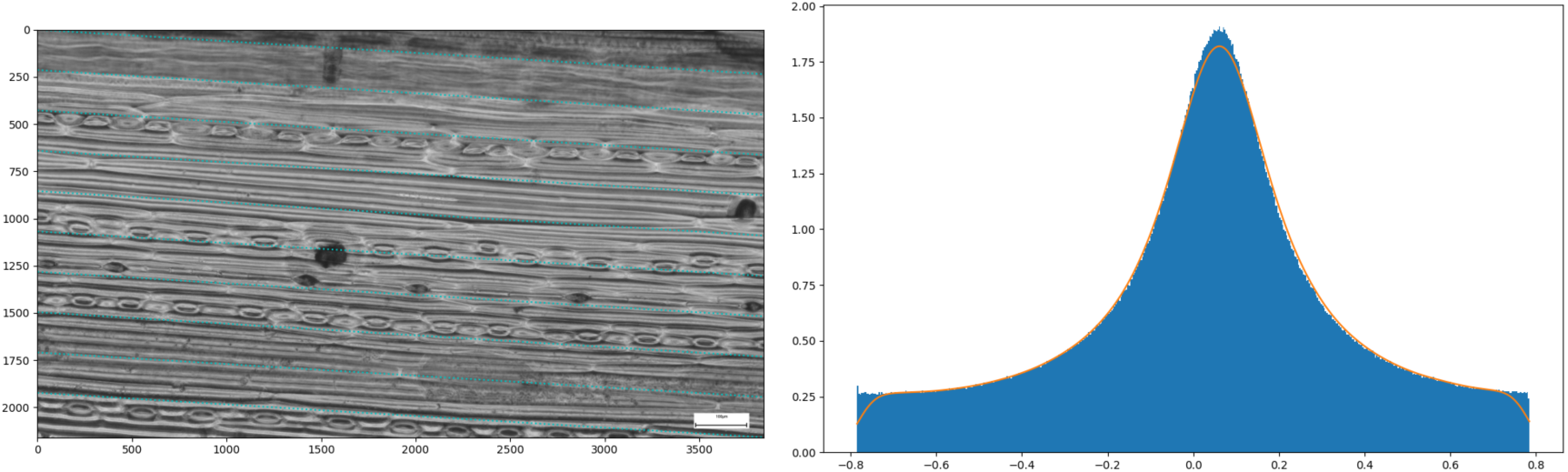
4. We used Cellpose 3.0 to perform cell segmentation on each image. Our goal is to automatically identify and segment one specific cell type (sister cells after differentiation) from each image. For this, we fine-tuned a pretrained model on hand-labeled examples. Since there is a large diversity of plant/cell types, in this study we train 12 different models (including 2 stomata cell models, see details below). To train each model, we need to provide examples of the cells we want to segment. We first crop an image so that it contains about 30 regions of interest (ROIs), then hand-label the ROIs using the Cellpose GUI interface. This is repeated at least 3 more times to obtain a training dataset. Using the images and masks from this dataset, we fine-tune the pretrained “cyto” model provided by Cellpose, with the greyscale channel of images, a learning rate of 0.04, weight decay of 0.0001, and 8 images per epoch. We choose the number of training epochs so that the training loss <0.3 at the end of training. Using this newly-trained model, we test on more images, identify poor segmentation results, then hand-label these failing examples (here, a filtration step is needed to remove tiny masks) to add to the training set. We repeat this procedure until the segmentation results are accurate. For regions where sister cells are not visible due to dense trichomes (in our data, the wheat elongation zone and the entire *Brachypodium* leaf), we instead identify trichomes as they grow alternatively with sister cells in the same cell file. Sister cell length can be inferred from adjacent trichome distances. For most species, a cell file of stomata is accompanied by two cell files of sister cells, thus the model is able to accurately identify the sister cells by their adjacent stomata. However, oat has two lines of stomata cells next to each other, and are then accompanied by two cell files of sister cells. Thus some cells adjacent to stomata in the stomata cell files cannot be distinguished from sister cells. To remove them, we identify stomata and filter out cells in the same cell files as stomata. The different models we trained are listed below:

a. Meristem cells in *Brachypodium* (1)
b. Meristem cells in Wheat (2), Oat (3), Barley (4)
c. Trichome identification and distance measurement in *Brachypodium* (5), Wheat (6)
d. Sister cells in Wheat (7) which mostly after cell elongation
e. Sister cells in Barley (8) which covers a large range of cell size
f. Smaller-sized sister cells(9) and young stomata cells in Oat (10)
g. Larger-sized sister cells(11) and mature stomata cells in Oat (12). Given the big cells, the images were shrunk in width to help finding intact cells
h. The images with poor cell segmentation results or very specific to samples can be hand-marked using Cellpose interface and fed into the next step directly.

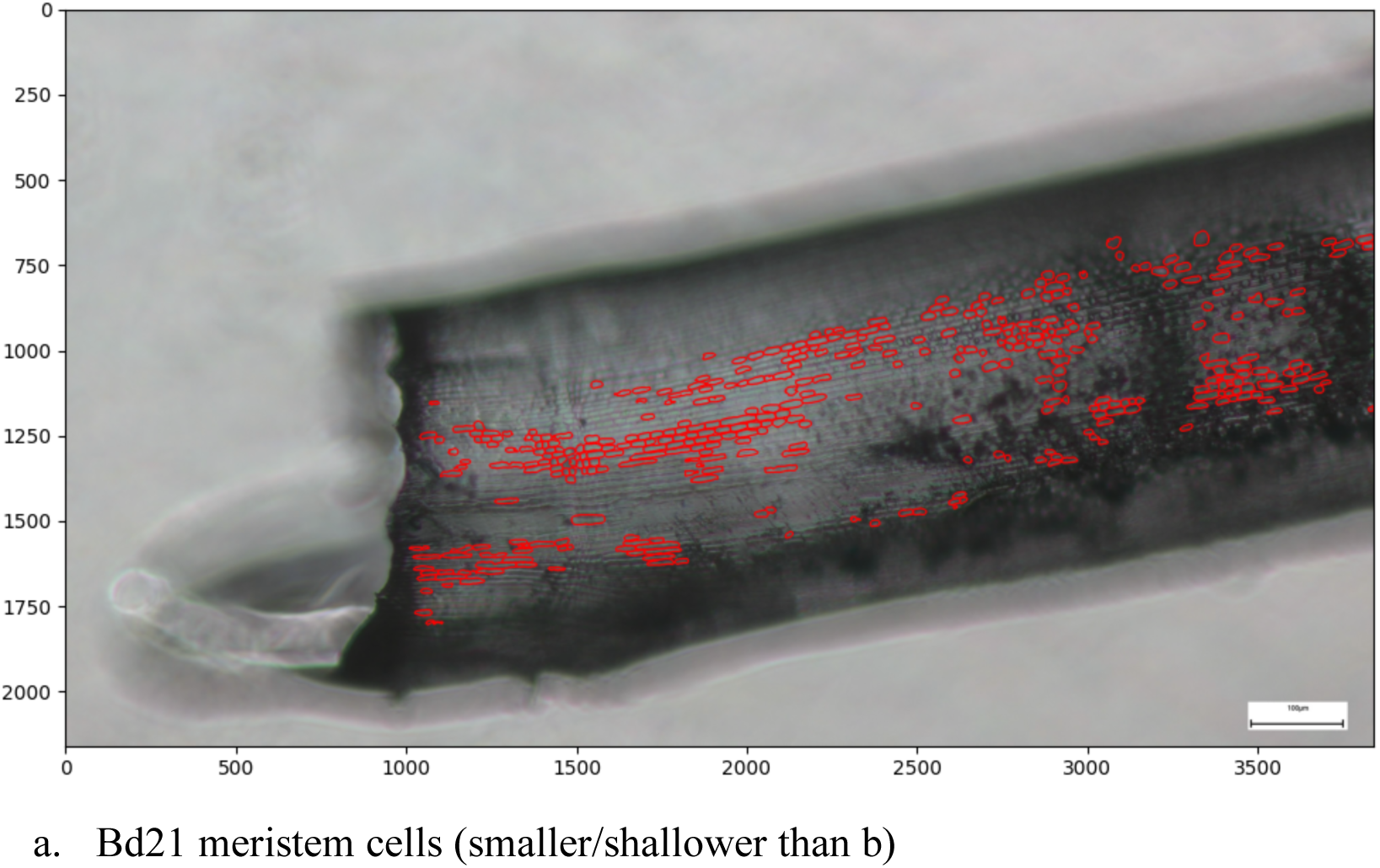
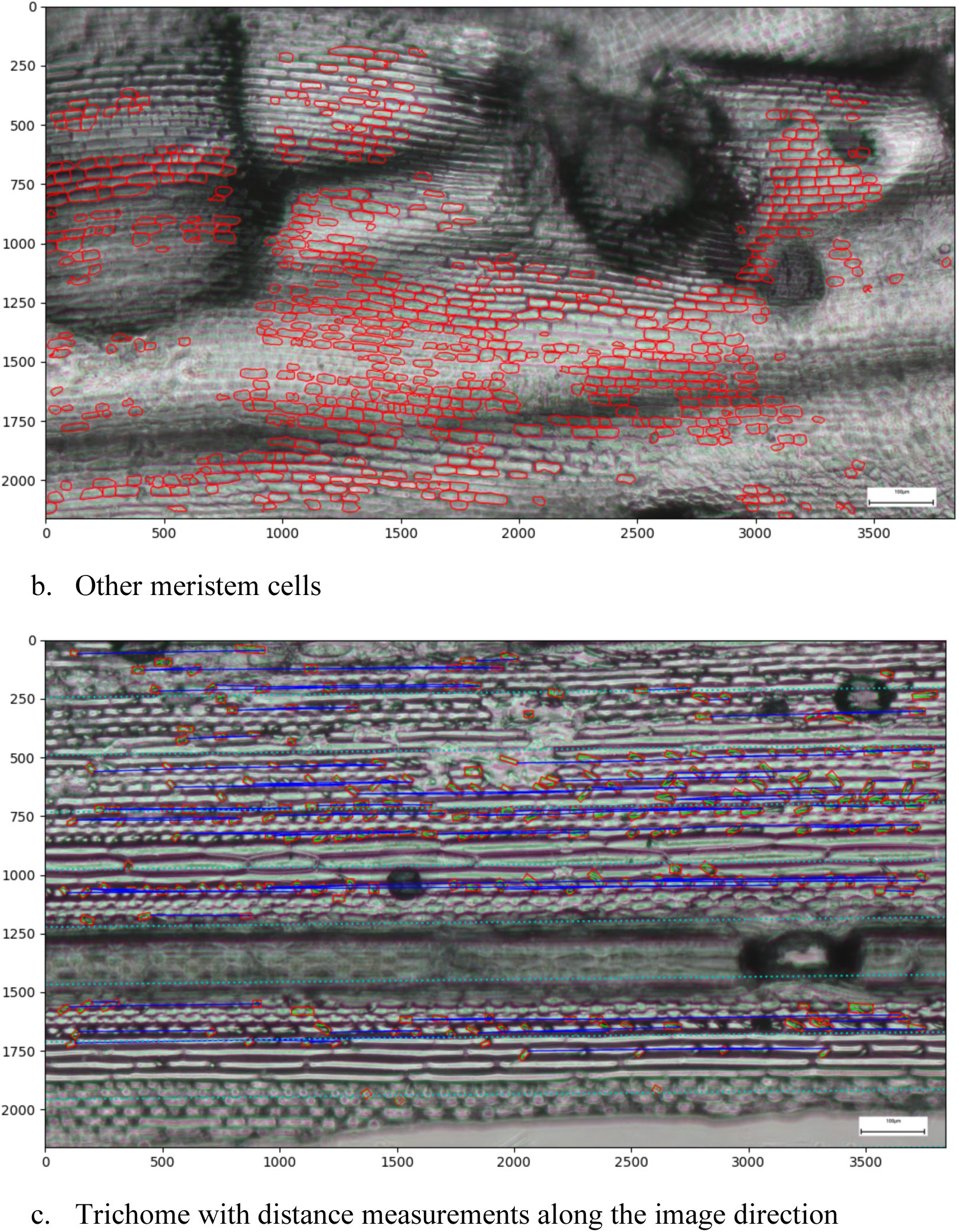
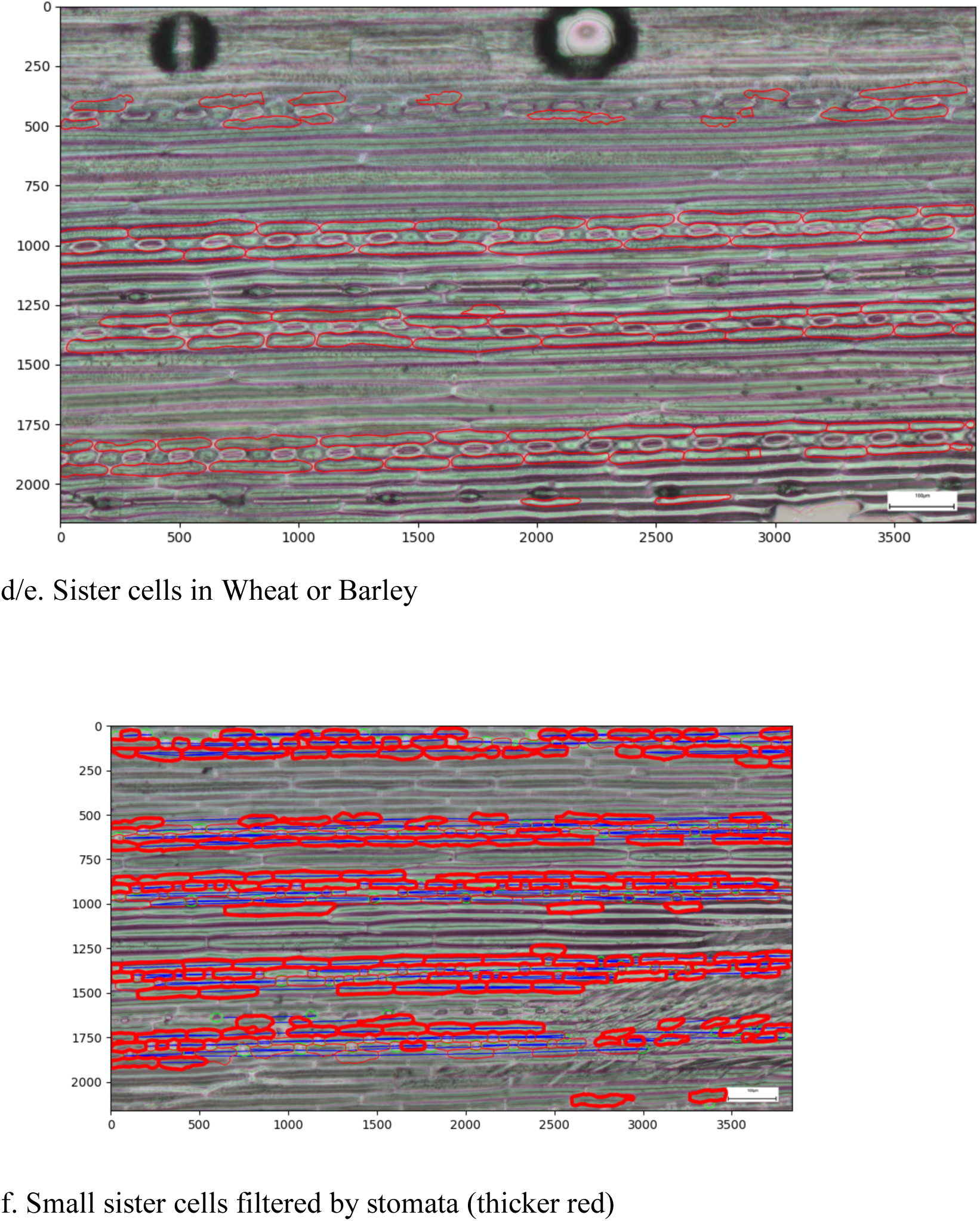
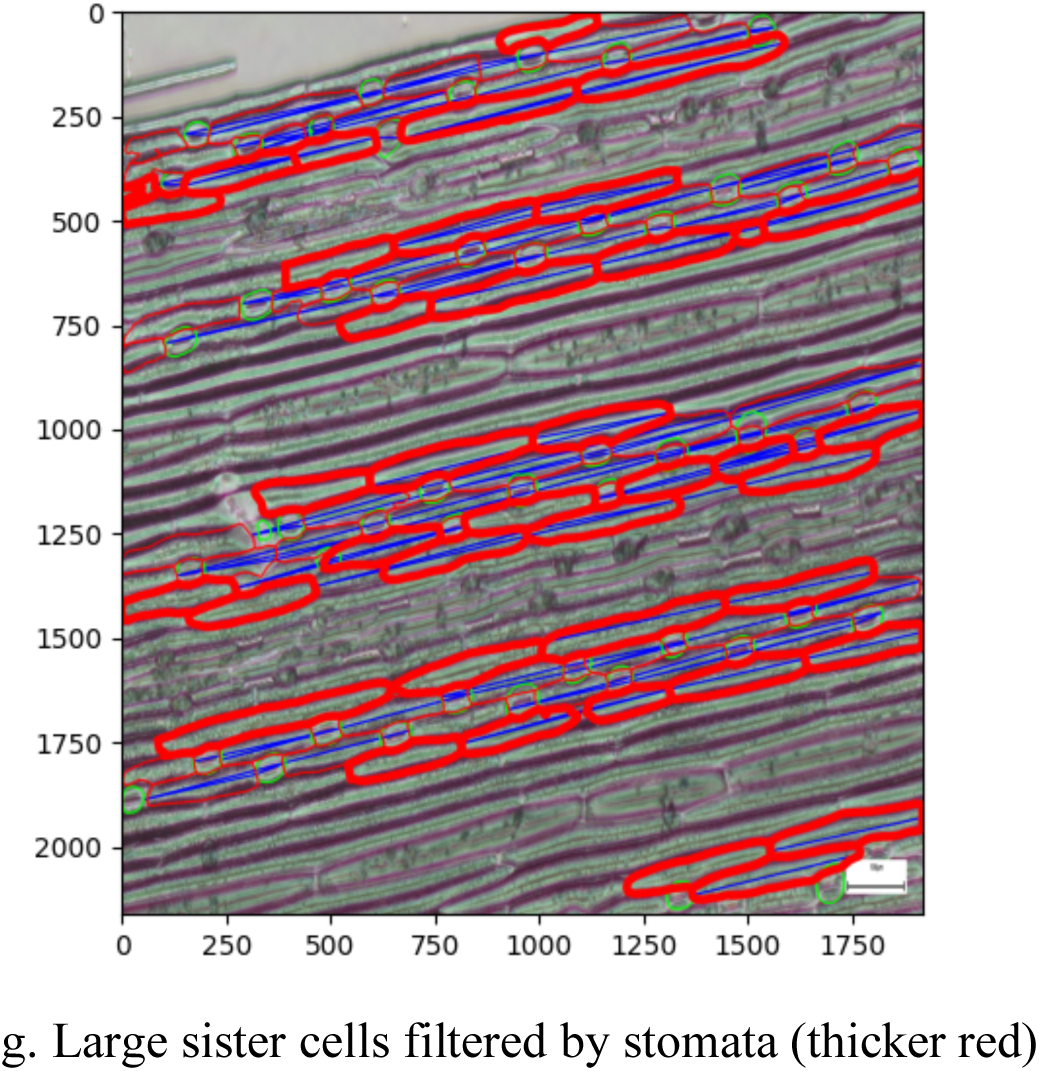
5. The cell segmentation results are then extracted with a custom script. There are three different post-processing steps depending on the cell type:

a. For meristem or sister cells, the ferit’s diameters (a box that can fit the cell) of the ROIs are identified together with the coordinates in the images, and they are saved into a .csv file.
b. For trichome cells, the distances between the closest two trichome cells are identified along the leaf directionality. The distance between the coordinates of the trichomes in the images are then used as a proxy for the diameter of the sister cell between the trichomes.
c. For oat sister cells, to remove cells that are in the same cell file as stomata cells, the stomata cells are first identified, then any cells belonging to the same cell file (inferred from the image directionality) as the stomata cells are removed. The sister cells left are then identified for cell size and coordinates.
6. The cell lengths and location along the leaf axis determined from the models described above are extracted with a custom R script. To filter outliers resulting from misidentification from the models, the cell sizes along the leaf axis are binned between 300um to 500um and the 50% quantile if number of cells in the bin >5 is used instead. The quantile of 30% for trichome distance with number of cells in the bin >20 are used. This results in a scatter plot of the cell length against leaf location for each leaf.
7. Overall although different models are needed to better capture the range of cell length in the images, we make sure there are overlap regions where the models have consistent cell length to avoid the effect of different models on the model fitting. For example, three models are shown in wheat.

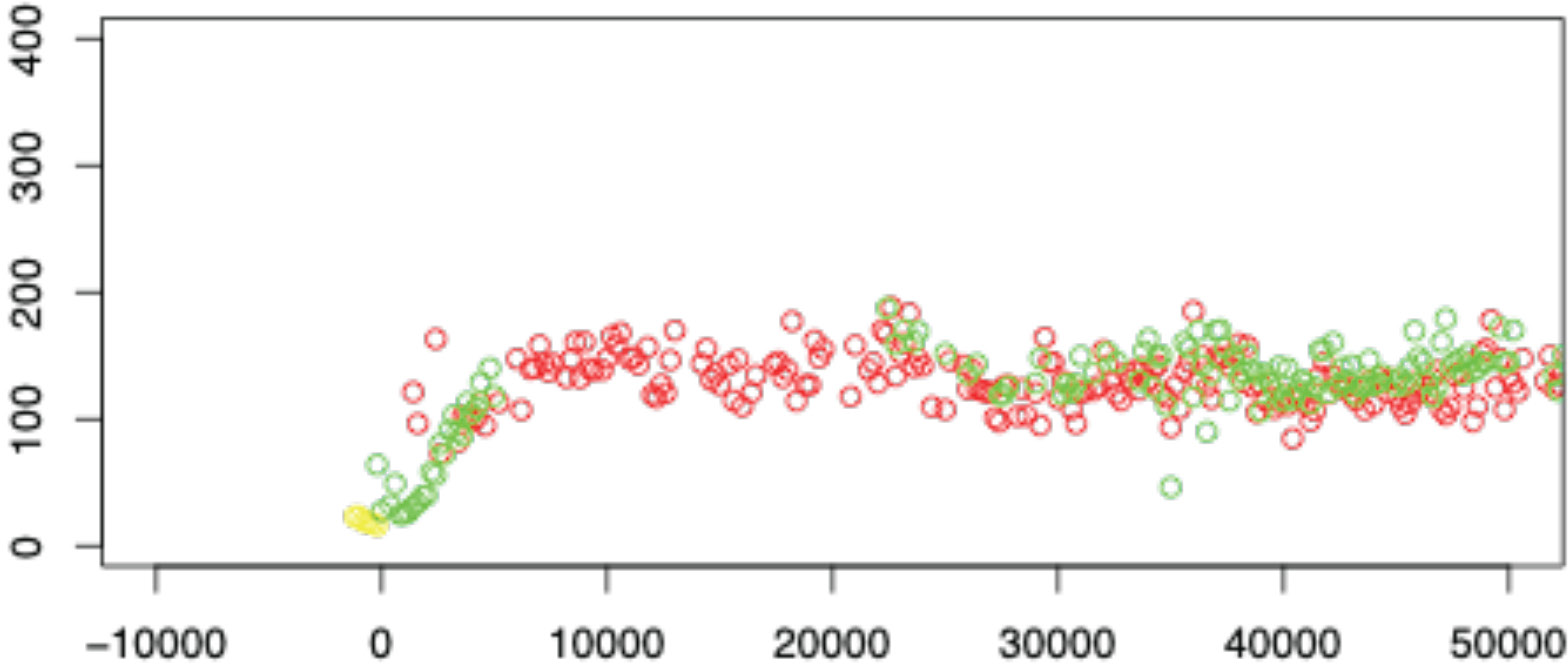
8. We then fit a sigmoid model to the scatter data of each leaf, using a custom python script. For samples with dropping tail, or quick dropping after elongation, or a dent in mature zone, we only fit the sigmoid model to before the dropping point.

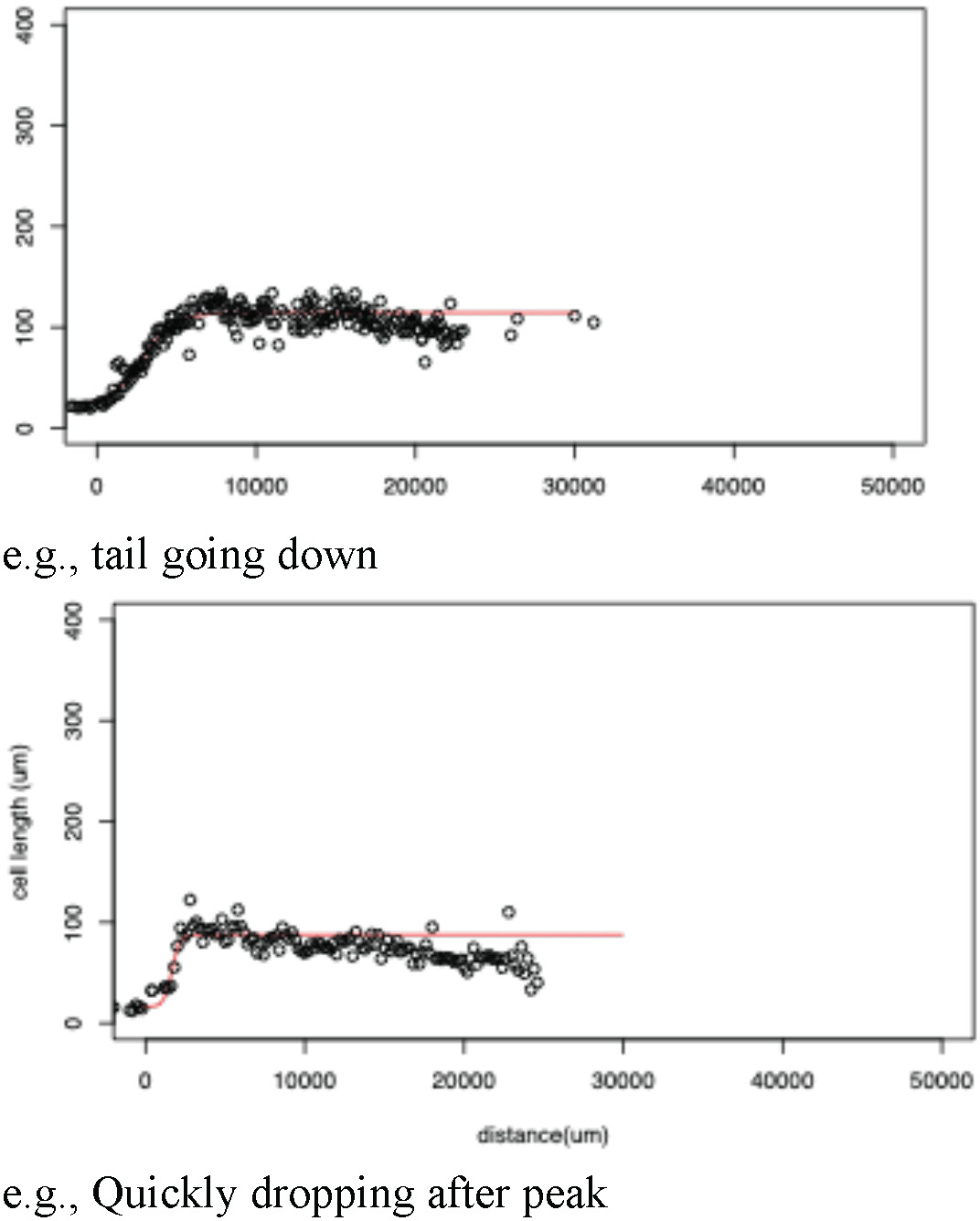
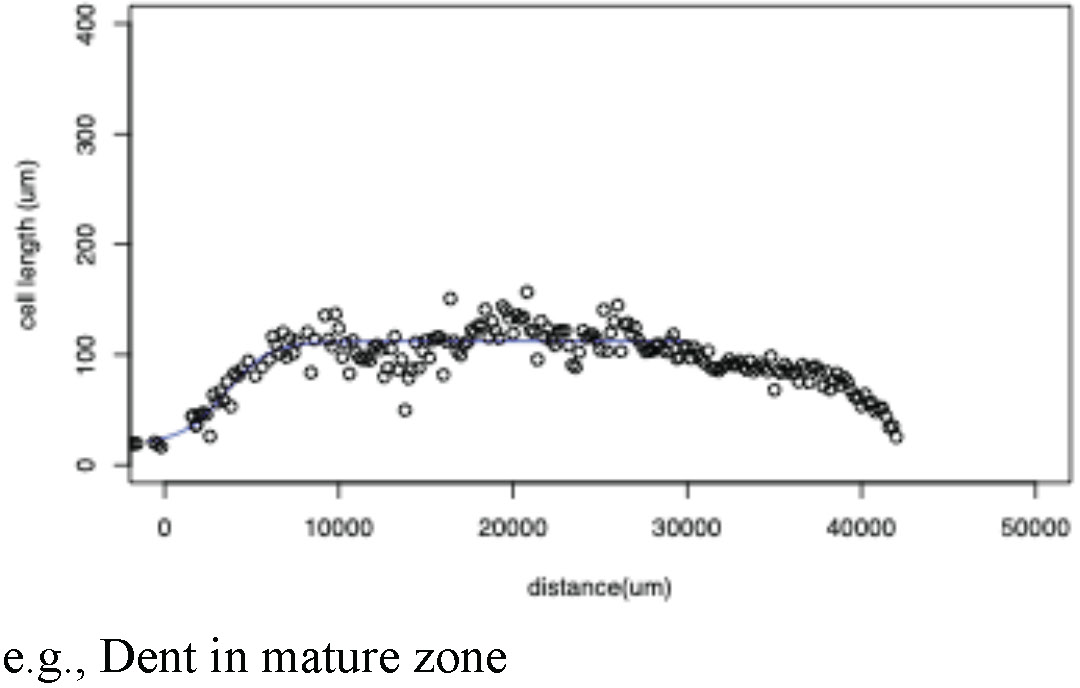

